# Identifying weak signals in inhomogeneous neuronal images for large-scale tracing of neurites

**DOI:** 10.1101/181867

**Authors:** Shiwei Li, Tingwei Quan, Hang Zhou, FangFang Yin, Anan Li, Ling Fu, Qingming Luo, Hui Gong, Shaoqun Zeng

## Abstract

Reconstructing neuronal morphology across different regions or even the whole brain is important in many areas of neuroscience research. Large-scale tracing of neurites constitutes the core of this type of reconstruction and has many challenges. One key challenge is how to identify a weak signal from an inhomogeneous background. Here, we addressed this problem by constructing an identification model. In this model, empirical observations made from neuronal images are summarized into rules, which are used to design feature vectors that display the differences between the foreground and background, and a support vector machine is used to learn these feature vectors. We embedded this identification model into a tool that we previously developed, SparseTracer, and termed this integration SparseTracer-Learned Feature Vector (ST-LFV). ST-LFV can trace neurites with extremely weak signals (signal-to-background-noise ratio <1.1) against an inhomogeneous background. By testing 12 sub-blocks extracted from a whole imaging dataset, ST-LFV can achieve an average recall rate of 0.99 and precision rate of 0.97, which is superior to that of SparseTracer (which has an average recall rate of 0.93 and average precision rate of 0.86), indicating that this method is well suited to weak signal identification. We applied ST-LFV to trace neurites from large-scale images (approximately 105 GB). During the tracing process, obtaining results equivalent to the ground truth required only one round of manual editing for ST-LFV compared to 20 rounds of manual editing for SparseTracer. This improvement in the level of automatic reconstruction indicates that ST-LFV has the potential to rapidly reconstruct sparsely distributed neurons at the scale of an entire brain.

## 1. Introduction

Neuronal morphology is usually considered the basic structural unit in the brain (Marx 2012). The reconstruction of neuronal morphology, including soma localization, soma shape reconstruction, neurite tracing, and spine detection, is a key issue in many areas of neuroscience research, such as identifying neuronal types, inferring structural connections between neurons, drawing neuronal circuits, and modeling neuronal functions (R. Parekh and Ascoli 2013; Donohue and Ascoli 2011; Lu 2011; Meijering 2010; Svoboda 2011). Neurites form the core of neuronal morphology (Ruchi Parekh and Ascoli 2015; Peng et al. 2015), and correspondingly, tracing neurites plays an important role in the reconstruction of neuronal morphology.

In recent years, a series of breakthroughs in molecular labeling (Feng et al. 2000; L. Luo and Callaway 2008; Ugolini 2010; Jefferis and Livet 2012) and optical imaging techniques (A. Li et al. 2010; Gong et al. 2013; Gong et al. 2016; Ragan et al. 2012; Silvestri et al. 2012; Osten and Margrie 2013; Chung and Deisseroth 2013) have enabled us to rapidly collect brain-wide neuronal images at submicron resolution. These imaging datasets display the detailed structure of neurons at an unprecedented scale and contain nearly complete morphological information about neurons (Osten and Margrie 2013). However, the high-throughput and accurate reconstruction of neuronal morphology from these large-scale datasets still faces many challenges. One key challenge is to automatically identify neurites with weak signals. This identification is universally required to reconstruct neuronal morphology from a large-scale dataset due to the complicated nature of neuronal morphologies and the characteristics of imaging datasets.

To some extent, neuronal morphologies are closely linked to the characteristics of imaging datasets. A certain level of weak signal exists, originating from the size differences in portions of neuronal morphology. Some soma radii can be up to several tens of micrometers, while the radii of some neurites are several hundred nanometers. When imaging neurites with small radii, a few fluorescent molecules can serve as labels, resulting in low signal intensity from these neurites. Images are collected at a relatively low spatial sampling rate, increasing the difficulty in identifying weak signals. Because neuronal morphology varies across different brain regions or even the whole brain, a balance must be achieved between the sampling rate and the imaging speed. Therefore, a relatively low sampling rate is practical. The background of large-scale neuronal images is also inhomogeneous, generally as a result of the complicated imaging procedure used and the structural differences in different brain regions. Some key characteristics of neuronal images can be illustrated in Fig. 1. Two sub-blocks were chosen from a whole-brain imaging dataset (Figs. 1a-c), both of which contained a neurite with weak signals whose signal-to-background-noise ratios (SBRs) are low, approximately 1.05 (Figs. 1d-f). Furthermore, as indicated in Fig. 1f, the background intensity of sub-block *b* is even higher than the foreground intensity of sub-block *c*.

**Figure 1.**
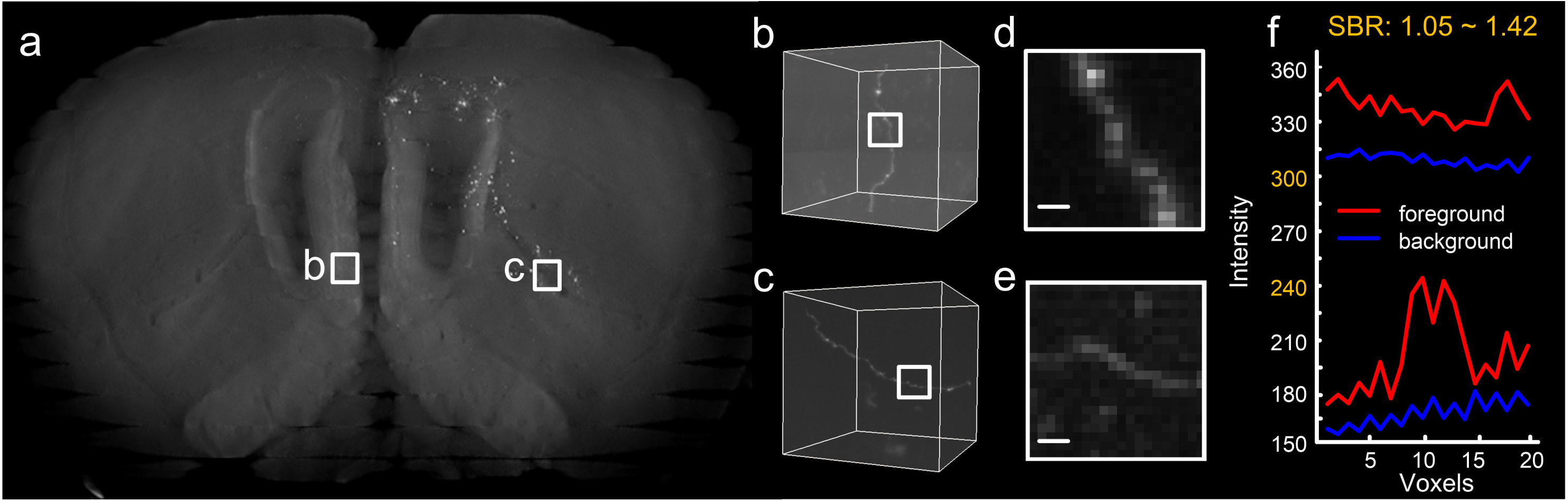
Some key characteristics of neuronal images at the brain-wide scale. (a) A thumbnail view of a mouse brain dataset in which two sub-blocks are highlighted with squares; (b) a sub-block with a single neurite, part of which is labeled with a square; (c) similar to (b); (d) maximum projections of the labeled view in (b) through a depth of 10 μm, with a scale bar of 2 μm; (e) similar to (d); (f) The upper two curves represent the foreground (red) and the background (blue) of the neurite in (d), and the bottom two curves correspond to the neurite in (e).

Many methods trace neuritis well and demonstrate a good ability to identify neurites with weak signals (De et al. 2016; Acciai et al. 2016; Sreetama Basu et al. 2016; Mukherjee et al. 2015; Xu and Prince 1998; Cai et al. 2006; Saurav Basu et al. 2013; Radojevic and Meijering 2017; Choromanska et al. 2012; Zhao et al. 2011; Santamaria-Pang et al. 2015; Wang et al. 2011; G. Luo et al. 2015; Peng et al. 2010; Turetken et al. 2011; Chothani et al. 2011; Yang et al. 2013; Bas and Erdogmus 2011; Rodriguez et al. 2009; Liu et al. 2016; T. Quan et al. 2016; S. Li et al. 2016; Frangi et al. 1998). Some of these methods are listed below: model-fitting (Zhao et al. 2011; Santamaria-Pang et al. 2015), open-snake (Wang et al. 2011; Cai et al. 2006; G. Luo et al. 2015; Xu and Prince 1998), graph-based (Peng et al. 2010; Turetken et al. 2011; Saurav Basu et al. 2013; Chothani et al. 2011; Yang et al. 2013), principal curve (Bas and Erdogmus 2011), voxel scooping (Rodriguez et al. 2009), multi-scale (Choromanska et al. 2012; Frangi et al. 1998), density filters (Radojevic and Meijering 2017), and others. However, most of these methods use a set of thresholds to determine whether the traced neurite continues or not, and thus may encounter difficulty separating weak signals from an inhomogeneous background. Recently proposed machine learning methods (R. Li et al. 2017; Chen et al. 2015; Megjhani et al. 2015; Gu et al. 2017; Hernandez-Herrera et al. 2014; Becker et al. 2013) can generate tracing results with better accuracy than traditional methods. However, these methods may fail to rapidly trace neurites in large-scale images. Machine learning methods consider many image features and use an algorithm to detect dominant image features, thereby requiring intensive computations. Some of these methods instead separate the identification and tracing procedures (R. Li et al. 2017; Chen et al. 2015; Megjhani et al. 2015) and attempt to identify as many signal voxels as possible, which generates the detailed shape of a neurite. However, heavier computation costs are incurred for larger images.

In this study, we propose a method for identifying weak signals and embed this method into the neurite tracing pipeline. Our strategy closely links the identification and tracing procedures, and requires only a few foreground voxels in the tracing process for identification. We observed many neuronal images and determined identification rules: the local background must be smooth, and the neurite must have a strong anisotropic shape that can be identified with a carefully chosen threshold value. These rules can be summarized as a feature vector that distinguishes foreground and background voxels. By training feature vectors on foreground and background voxels, we obtained a classifier (Suykens and Vandewalle 1999; Cortes and Vapnik 1995) that we combined into our previous tool, SparseTracer (S. Li et al. 2016), naming the combination tool SparseTracer-learned feature vector, ST-LFV. We verified that ST-LFV can identify weak signals well from relatively low-sampling-rate images and can overcome the identification difficulties caused by inhomogeneous backgrounds. We demonstrated that ST-LFV significantly enhances the performance of SparseTracer in large-scale tracing of neurites.

## 2. Methods

In this section, the components of ST-LFV are outlined. First, we describe the method used to extract the feature vector of a voxel, which displays the differences between the foreground and background voxels. Second, we introduced support vector machine (SVM) to the feature vectors space to build an identification model that can detect weak signals (Suykens and Vandewalle 1999; Cortes and Vapnik 1995). Third, we implemented this identification model as part of our previous work, SparseTracer (S. Li et al. 2016), for the purpose of neurite tracing. We also discuss how to select parameters in ST-LFV.

### 2.1 Feature extraction for identifying weak signals

The extraction of representative image features is key to identifying weak signals and is based on premises drawn from the images themselves. Here, our premises are that the shape of a neurite can be described by a series of touching cylinders; the background is locally smooth; and, in a small local region, the foreground and background intensity can be distinguished with a well-chosen threshold value.

In general, for a given voxel, the extraction of its corresponding features consists of several steps: a) set a series of threshold values, in descending order; b) for each threshold value, generate the growth region of the current voxel; c) form a feature vector for this voxel composed of elements corresponding to the rate at which regional volumes grow to fill the pre-determined neighborhood volume of this voxel, called the volume filling rate. Note that the pre-determined neighborhood used in this study contained 19 × 19 × 19 voxels whose size is much larger than the radius of a neurite. We describe how to extract features in detail and explain why the extracted features are consistent with our premises.

First, set a series of threshold values to generate the growth region. For a given voxel, we collected in its neighborhood a number of voxels that connect with each other and whose signal intensities are greater than the pre-determined threshold. These collected voxels form a growth region. In obtaining a growth region, the threshold setting is a key factor. Here, we set a series of descending threshold values for a given point *p*^*^ in an image:

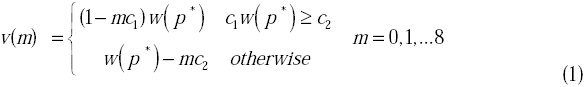

Here, *p*^*^ represents a 3-dimensional vector containing *x, y*, and *z* coordinates; the rounding operation of *p*^*^ equals its corresponding voxel *o*^*^; and *c*_*1*_ and *c*_*2*_ are two predetermined constants, *c*_*1*_ = 0.025 and *c*_*2*_=1.5. Here, 1-m*c*_*1*_ is a ratio. *w*(*p*^*^) is a weighted value calculated from the signal intensities and the positions of the voxels adjacent to *o*^*^, and provides a robustness result. This value can be calculated as

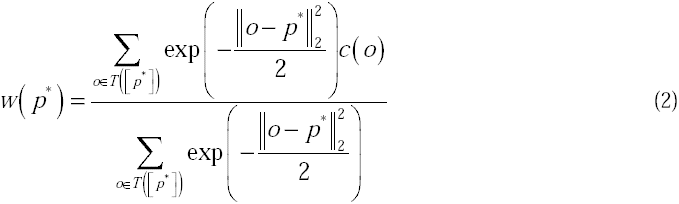

Here, *T* is the set that includes [*p*^*^] and its 6-voxel neighborhood, [] represents the rounding operation of the point’s coordinates; we denote the voxel *o*^*^ by [*p*^*^]; *c*(*o*) is the intensity value of the voxel *o*; and || ||_2_ represents the 2-norm.

The threshold value is codetermined by a series of ratios and the weight value of a given point. The threshold values vary when the weighted value changes. To simplify the form of this expression, threshold values are represented by invariable ratios, denoted as 1-*mc*_1_, where *m*=0,1,…,8.

When the signal value of a given point is small, the decrease in the threshold values *v*(.) calculated by Eq. 1 varies extremely slowly. To avoid this case, we set a lower bound (*c*_*2*_ = 1.5) to meet the decreased amplitude of the threshold values.

We next generate the growth region with a pre-determined threshold value. For the given voxel *o*^*^, equal to [*p*^*^], the growth region is generated in its neighborhood region *N*, which is a cubic region of 19 × 19 × 19 voxels with *o*^*^ as its central voxel. The steps used to generate the growth region are listed below.

a) Set the initial seed as the voxel *o*^*^, labeled with an arrow in Fig. 2a, and search for its neighboring voxels whose intensity *c*(*o*)satisfies Eq. 3

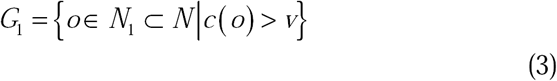

**Figure 2.**
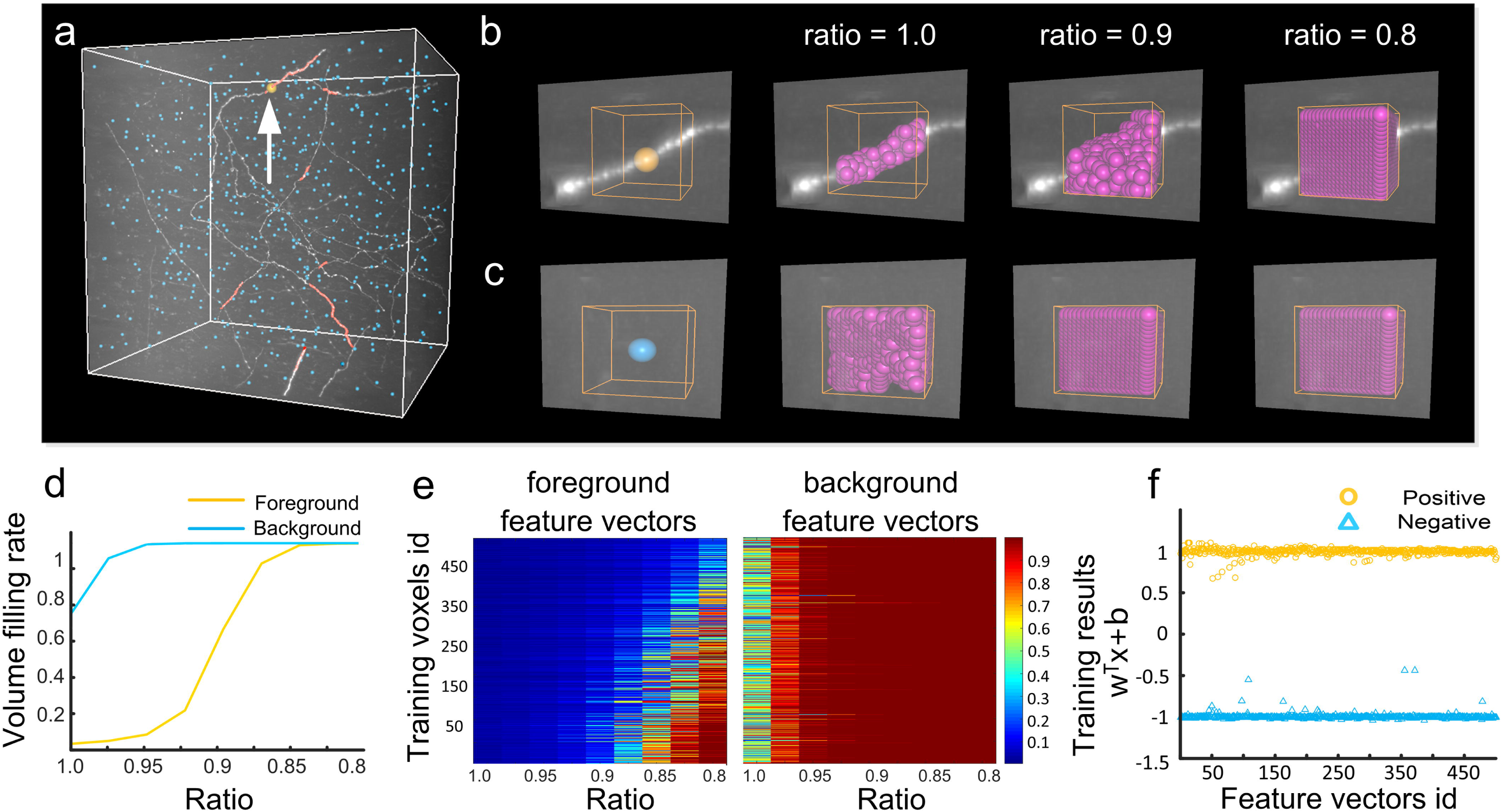
An illustration of feature vector extraction. (a) An image dataset including the labeled foreground voxels (light red) and background voxels (light blue). The foreground voxel (orange) labeled with an arrow and its corresponding growth regions in (b). (b) Calculating the growth regions of the selected foreground voxel (yellow) respective to different ratios. The neighborhood and the growth regions of this voxel are labeled by the yellow cubic and purple points, respectively. (c) The same as (b) for the selected background voxel (blue). (d) The foreground (yellow line) and background (blue line) feature vectors calculated from the given voxels in (b) and (c), respectively; (e) feature vectors of all labeled foreground (left) and background (right) voxels in (a); (f) generating an SVM identification model with feature vectors from (e). The positive (yellow circles) and the negative (blue triangles) results corresponding to the foreground and background feature vectors in (e), respectively.

Here, *N*_1_ is the 26-voxel neighborhood of the voxel *o*^*^; *v* is one of the threshold values calculated by Eq. 1; and the voxel *o*^*^ and the searched voxels form the voxel set *G*_1_, which is then labeled.

b) In the unlabeled region of *N*, search for the 26-voxel neighborhoods of every voxel in the set *G*_1_, denoted by *N*_2_. According to *N*_2_ and the threshold *v*, use Eq. 3 to generate *G*_2_, and then label *G*_2_.

c) Repeat the above procedure until no new voxel sets can be generated in *N*. All labeled voxels form the growth region of the voxel *o*^*^ with respect to the threshold *v*, denoted by *G*(*v*).

The above procedure used to generate a growth region is similar to that described in our previous work (S. Li et al. 2016), and a series of growth regions can be generated when the threshold value in Eq. 1 varies. Figs. 2b and c illustrate the procedure of obtaining the growth region of a signal voxel and a background voxel, respectively. The total number of the labeled voxels is determined by the current threshold. Note that according to the specific expression of the threshold in Eq. 1, the same ratios do not represent the same threshold values for different voxels.

Next, calculate the feature vector of a given voxel. For a given voxel, we can obtain its corresponding growth region with a pre-determined threshold. The volume filling rate between the growth region volume (number of labeled voxels) and the neighborhood region volume (total number of voxels in the neighborhood) is then computed. The feature vector of a point is composed of a series of rates derived from different ratios, and this vector is denoted by *R*. The *m*^th^ component of *R* is defined by

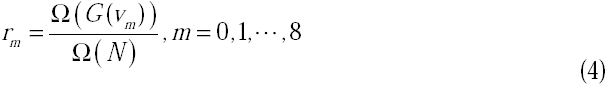

where Ω(□) is the total number of voxels in the set.

Following the above mentioned steps, we can achieve the corresponding feature vector for every voxel (both foreground and background) in the image.

Here, we explain why the extracted features are consistent with our premises. If a point belongs to the background, the volume filling rate in the feature vector will increase to 1.0 rapidly because of the smoothness of the local background. For the feature vector of a foreground point, the volume filling rate will increase much more slowly, and may not even reach 1.0, as the cylindrical shape of a neurite takes up a small amount of space in its neighborhood and the intensities of foreground and background voxels will vary. The differences between foreground and background feature vectors are shown in Fig. 2d.

### 2.2 The SVM identification model used to identify weak signals

This section is composed of two parts: 1) automatically extracting raining sets from neuronal images; 2) generating the identification model after obtaining the training set.

In a supervised learning framework, a training set is necessary. Here, the training set used contains the feature vectors of foreground and the background points. The automatic generation of training sets thus requires only computationally obtaining some foreground and background points, which may be practical for the following reasons. Existing tracing methods can identify weak signals at a certain level, which can provide foreground points. The voxels of neurites comprise an extremely small proportion of the total voxels in neuronal images. This indicates that there is an extremely small chance that some foreground points are included in the several hundred background points that are randomly chosen (uniformly distributed) from neuronal images.

Here, we used our previous tool, SparseTracer, to trace neurites and extract foreground points from the traced results. The traced results provide the skeleton of a neurite, being composed of a series of points in which the adjacent points connect. These skeleton points can be recognized as foreground points (Chen et al. 2015). If fewer than 500 skeleton points are selected, we calculate the feature vectors of all skeleton points. Otherwise, we choose those skeleton points with median signal intensities and calculate their corresponding feature vectors. These feature vectors constitute the positive training set, denoted by *S*={*s*_1_,*s*_2_,*…,s*_*n*_}, (Left, Fig. 2e).

To obtain negative training samples, we randomly (from a uniform distribution) selected points from neuronal images that had same number of positive training samples and calculated their feature vectors, denoted by *B*={*b*_1_,*b*_2_,*…,b*_*n*_} (Right, Fig. 2e). Considering that the number of foreground voxels is extremely small compared to the total number of voxels in neuronal images, the probability that a selected point is a foreground point is extremely low. Furthermore, the positive training set is known. We identified whether a feature vector in *B* was an outlier by measuring two degrees of similarity: one is the inner product between this feature vector and the mean values of the negative training set *B*, and the other is the inner product between this feature vector and the mean values of the positive training set *S*. If the former value is larger than the latter, the vector is regarded as an outlier and is deleted from the dataset *B*. The remaining vectors in the dataset comprise the negative training samples.

To simplify descriptions, we used {*y*_*k*_, *x*_*k*_},*k=*1,2, …, *K* to denote the positive and the negative training sets. Here, *y*_*k*_=1 or −1, *x*_*k*_ is a feature vector, and *K* is equal to the sum of the number of training feature vectors in the set *S* and in the set *B*. If *y*_*k*_=1, *x*_*k*_ is then positive and equal to a component in the set *S*. Otherwise, *x*_*k*_ is equal to a component in the set *B*. After obtaining the training set, we introduced a support vector machine (SVM) (Suykens and Vandewalle 1999; Cortes and Vapnik 1995) to build a supervised classifier that distinguishes between foreground and background voxels. Constructing a supervised classifier can be equivalent to searching an optimal hyperplane that separates the positive training and negative training samples with maximum margin criteria. The optimal hyperplane can be mathematically described by

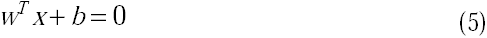

This hyperplane was determined by solving the classifying problem [49] as

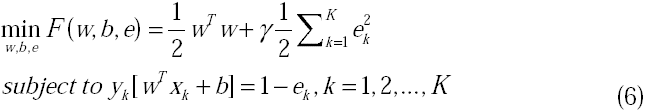

Here, 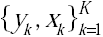 are training samples; as described above, *x* refers to the *k*^*th*^ feature vector. If *y*_*k*_ = 1, *x*_*k*_ is positive and is negative otherwise. ^γ^ is used to control the tradeoff between training error and generalization ability. The corresponding Lagrange problem is given as

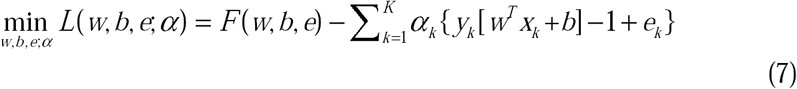

Where *α*_*k*_ are the Lagrange multipliers. Using Kuhn-Tucker conditions, we can obtain an optimal solution (Suykens and Vandewalle 1999), and the optimal hyperplane is given by

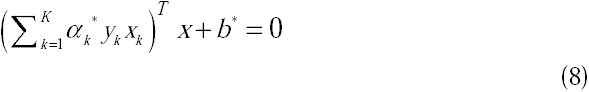

The corresponding supervised classier can be denoted by

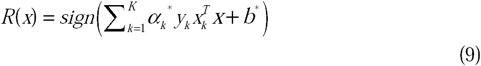

We used this classifier to identify whether a point belongs to the foreground, using the following method. Submit the feature vector of a point into the classifier. If the resulting value is greater than zero, the point is a foreground point; otherwise, it is a background point. We also applied the classifier to the training samples. The results show that most of the positive and negative values are nearly 1 and −1, respectively (Fig. 2f), which illustrates the large differences in the feature vectors of foreground and background points (Fig. 2e).

### 2.3 Using the identification model for neurite tracing

Neurite tracing is the process of obtaining the skeleton of a neurite. The skeleton is formed from tracing points in which any two adjacent points connect. When tracing a neurite, if the current tracing point is identified as a background point, the tracing will be terminated. Therefore, accurately identifying foreground points is a key component of neurite tracing. We applied the developed identification model to our previous tool, SparseTracer, to obtain better neurite tracing results. Specifically, the pipeline is described as follows and is shown in Fig. 3.

**Figure 3.**
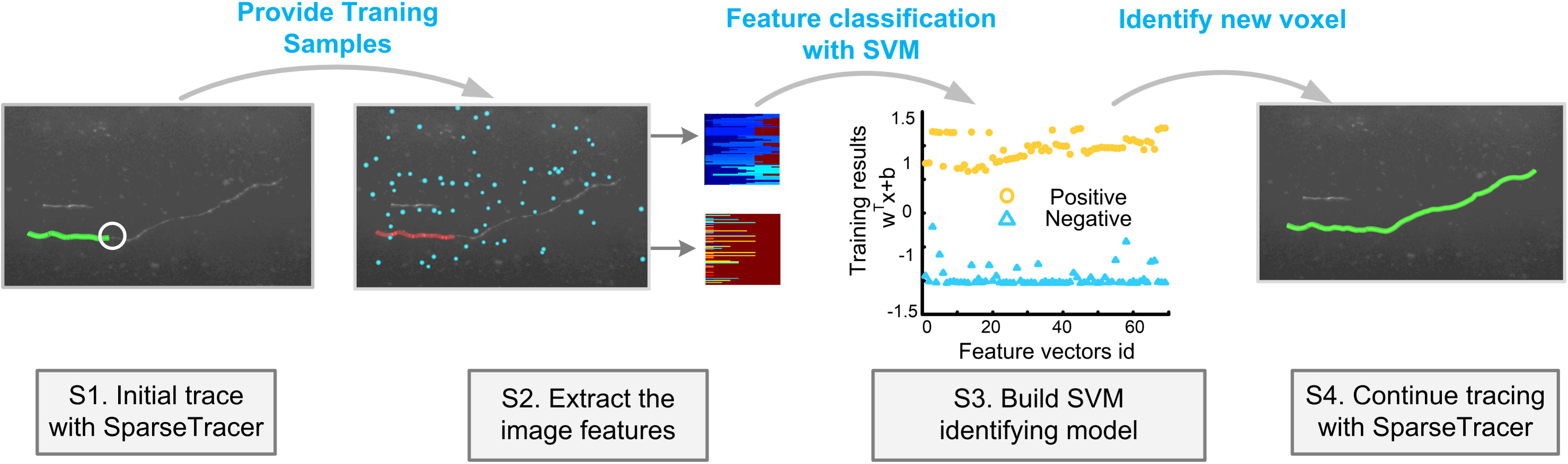
The pipeline of neurite tracing with ST-LFV. S1. Acquire the initial skeleton (green curve) of the neurite with SparseTracer. The site where SparseTracer fails is labeled with a circle; S2. Extract the feature vectors from the labeled voxels. These labeled voxels include foreground voxels (red) on the initial skeleton in S1 and background voxels (blue), and their corresponding feature vectors are located in the upper and lower panels, respectively; S3. Construct an SVM identification model with feature vectors in S2; S4. Use the identification model to identify weak signals and continue tracing. The final reconstruction (green) is thus obtained.

**Step 1)** Use SparseTracer to trace the neurite. When the point *p*_*n*+1_ is identified as a background point, tracing stops and an initial skeleton is generated, represented by *P* = {*p*_1_, *p*_2_,…, *p*_i_,…, *p*_n_}. Here *p*_i_ is the *i*^th^ point on the skeleton.

**Step 2)** Extract the feature vectors of foreground and background points separately; these form the positive and negative training sets, respectively. Note that the foreground points are the point set *P*.

**Step 3)** Obtain the SVM identification model with the training set.

**Step 4)** Apply the obtained model to the identification of the points *p*_*n*_ and *p*_*n*+1_. If one of these two points is identified as a foreground point, continue tracing with SparseTracer. Otherwise, terminate tracing.

Our identification model can be combined with most existing tracing methods to improve the process of neurite tracing. This statement is based on the fact that most tracing methods require identifying whether the current tracing point is a foreground or background point. Here, we combined our identification model with SparseTracer to improve tracing performance. SparseTracer can successfully trace a neurite with a relatively weak signal, which provides as many positive training samples as possible. In addition, SparseTracer can rapidly trace a neurite at a large scale, which is important for the complete reconstruction of a neuron.

When analyzing images, the SVM identification model can be updated iteratively to detect weaker signals. During neurite tracing, some signal points are identified by the initial model, and their corresponding feature vectors are added into the positive training set. The updated training set then generates the updated identification model. The updated model can detect weaker signals, as the added positive samples provide information on weaker signal points. Repeating the above procedure, more neurites can be traced, and we used this strategy in our analysis.

### 2.4. Parameter settings in the identification model

To construct the SVM identification model to detect weak signals, certain key parameters must be pre-determined, including the size of the training set, the ratios used in feature vector extraction, and the size of the neighborhood.

**The size of the training set:** The positive training set depends on tracing results. If the total number of foreground points in the traced neurites is less than the pre-determined number (500), we select all foreground points and calculate their feature vectors to form the positive training set. In this case, though the size of the positive training set is small (dozens of points), the identification model can still behave well, and therefore we did not use upsampling to increase the size of the training set. Otherwise, when the traced neurites include many points, many positive feature vectors can be generated. In this case, the upper limit for the number of feature vectors in the positive training set is 500. The selection of this number is based on a tradeoff between computation cost and classification performance. More training samples may not improve identification performance. This selection can ensure that the time required to identify weak signals is approximately the same as that required for neurite tracing. In addition, balanced training sets are helpful for a supervised SVM classifier, and the negative training set is therefore nearly the same size as the positive training set (Tang et al. 2009).

**The ratios used in feature vector extraction:** The feature vector of a point depends on the setting of ratios, as described in Eq. 1. In our analysis, the ratios used in descending order range from 1 to 0.8, and the difference between two adjacent ratios is *c*_1_*=*0.025. The choice of a small *c*_1_ is based on one of our premises, namely, that the local background is smooth. Consequently, a slight decrease in the ratio used (i.e., a small *c*_*1*_) can fill the entire neighborhood of a background point through region growth, and the corresponding component of its feature vector will be equal to 1. Therefore, a small value of *c*_1_ can capture the smoothness of the background. A rapidly decreasing ratio (i.e., a large *c*_1_) indicates lower threshold values. With these lower thresholds, the growth region of a weak signal point will quickly fill its entire neighborhood, and thus the features of the neurite’s morphology will not be captured. Overall, the selection of a small c_1_ is intended to capture the feature differences between weak signal points and background points.

**The size of the neighborhood in feature vector extraction:** In feature vector extraction, the neighborhood of a point contains 19 × 19 × 19 voxels. This is based on the following considerations: if the size of a neighborhood is small, the local morphology of a neurite extracted with a relatively low threshold may span the entire neighborhood in some situations, preventing the capture of its local morphology. However, a large neighborhood gives rise to the need for highly complex computation to obtain the growth region, which is a key step in feature vector extraction. Taking into account the local shape of a neurite whose radius is the size of approximately 1-3 voxels, we set the size of a neighborhood at 19 × 19 × 19 voxels. This setting satisfies the condition that the local morphology of a neurite occupy a small portion of the neighborhood.

All of the parameters discussed here remain unchanged in our analysis.

## 3. Results

The experimental datasets used in our analysis include the fMOST (Gong et al. 2013), DIADEM (Brown et al. 2011) and BigNeuron datasets (Peng et al. 2015). The FMOST datasets include typical sub-blocks from an imaging dataset of a single mouse’s whole brain. This whole-brain image dataset was collected with the fMOST imaging system (Xiong et al. 2014), with a voxel size of 0.3 × 0.3 × 1 um^3^. The DIADEM (http://www.diademchallenge.org) and BigNeuron (http://alleninstitute.org/bigneuron/data/) datasets are freely available; information about these datasets can be found at the abovementioned websites, respectively. We performed experiments on a computer workstation (Intel® Xeon® CPU 3.46 GHz computing platform, Quadro K4000 3G GPU, 192 GB RAM, Windows 7). Our analysis involved two algorithms: an automatic tracing algorithm, SparseTracer, and a combination of SparseTracer and the learned feature vector (LFV), ST-LFV. When using SparseTracer to analyze each image stack, we carefully chose the parameters used to produce better tracing results. When using ST-LFV, the default settings for tracing parameters were used, which can provide an initial training set in most situations; the identification parameters are fixed in the following analysis and have been discussed in detail in the Methods section.

We demonstrated that ST-LFV can trace neurites in inhomogeneous neuronal images. We selected two image stacks and the related image features are displayed in Figs. 4a-4f. One dataset had a strong background (Fig. 4a) and the other had a weak background (Fig. 4d). By enlarging the selected sub-blocks (Figs. 4b and c), differences are visible between the foreground and the background intensities. These differences provide a basis by which a human annotator can obtain the ground truth of the neurites (Examples: red curves in Figs. 4a and d). We estimated the background values of the voxels labeled with red curves (Figs. 4a and d) using our previous method (S. Li et al. 2016) and calculated the foreground values of the voxels with Eq. 2. In most cases, the background intensities (blue curve in Fig. 4c) estimated from Fig. 4a are 2-3 times larger than the foreground intensities (red curve in Fig. 4c) calculated from Fig. 4c. This result indicates that background intensities vary sharply in brain-wide imaging datasets. In this case, we compared the tracing results drawn from SpareTracer and ST-LFV (Figs. 4g and h, respectively). SparseTracer failed to trace one neurite (red curve in Fig. 4a) regardless of the tracing thresholds used, and successfully traced the other neurite with a well-chosen threshold. ST-LFV can provide trace results nearly equal to the ground truth (red curves in Figs. 4a and d). We conclude that ST-LFV can overcome the influence of varying background intensity on tracing results. Note that for SparseTracer, ‘high threshold’ refers to the default threshold set in the algorithm, which can provide robust tracing results in most cases; ‘low threshold’ refers to a well-chosen threshold that produces better tracing results in a specific case.

**Figure 4.**
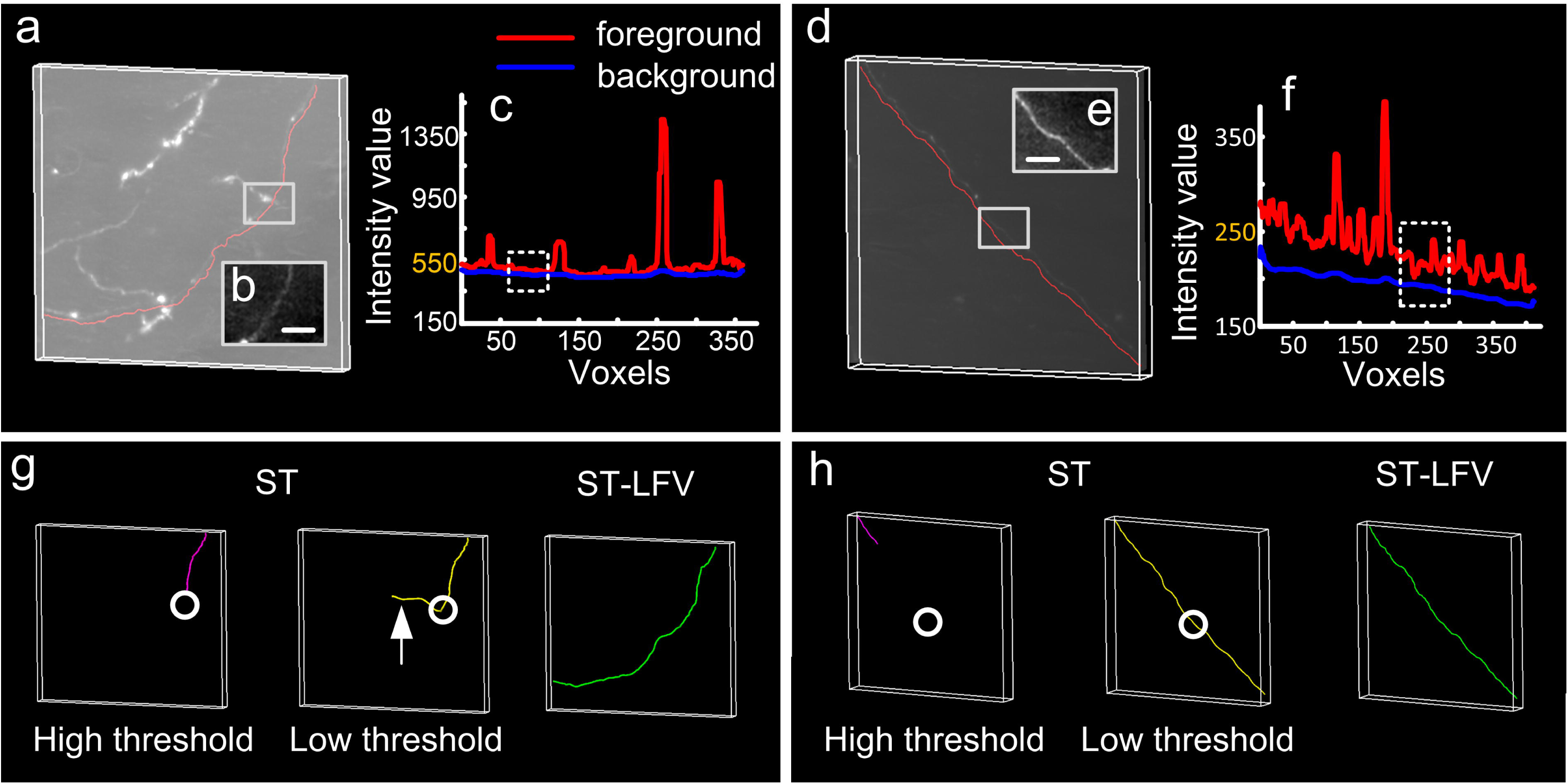
Performance on datasets with inhomogeneous backgrounds drawn from SparseTracer (ST) and ST-LFV, respectively. (a) One dataset with a strong background and a manually traced neurite (red). Part of this traced neurite is labeled with a square. (b) Maximum projections of labeled regions in (a) through a depth of 10 μm, with a scale bar of 10 μm; (c) The foreground (red) and background (blue) intensities of the traced neurite in (a). The intensities of the neurite in (b) are close to its background, labeled with dashed square; (d) One dataset with a weak background and one similar to (a); (e) and (f) have similar descriptions as (b) and (c), respectively; (g) Tracing results drawn from ST with a high threshold (purple) and a low threshold (yellow), respectively, and drawn from ST-LFV (green). The location of the weak neurite in (b) is labeled with circles. Over-traced results (arrow) drawn from ST with a low threshold; (h) similar to (g).

We used an experimental dataset to verify that ST-LFV can identify weak signals. The experimental dataset used includes a long neurite with weak signals at several sites (Fig. 5a). We selected two sites and extracted their corresponding sub-blocks (Figs. 5b and c). With the same procedure used to produce Figs. 4c and f, we estimated the foreground and background values of voxels from the two sub-neurites (Figs. 5b and c). This estimation indicated that these two sub-neurites have extremely weak signals and signal-to-background ratios (SBRs) as low as 1.05 (Figs. 5d and e). SparseTracer cannot address this case and only traces part of this neurite (Fig. 5f), despite our best efforts to select threshold values. Compared to SparseTracer, ST-LFV can produce perfect tracings (red curves, right in Fig. 5f). ‘High threshold’ and ‘low threshold’ in Fig. 5f have the same meanings as in Figs. 4g and h.

**Figure 5.**
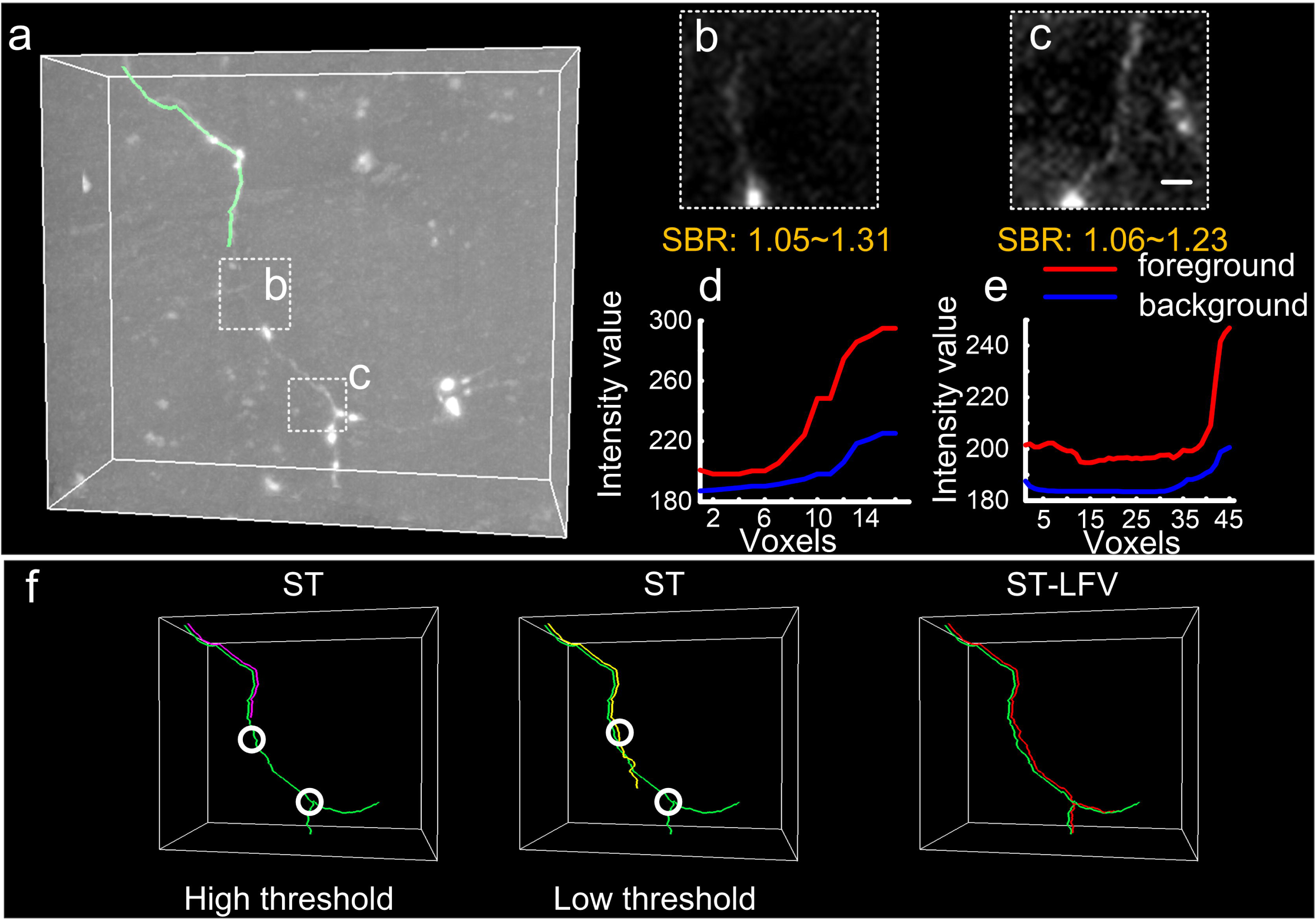
Performance on a dataset with weak neurites drawn from SparseTracer (ST) and ST-LFV, respectively. (a) A neurite and its initial tracing skeleton (green) drawn from ST. Two typical weak signal regions (dashed squares b and c) contain portions of this neurite; (b) and (c) are the enlarged views of the labeled regions in (a). Both include maximum intensity projections through a depth of 10 μm, with a scale bar of 5μm; (d) and (e) are the foreground and background intensities of the neurites in (b) and (c), respectively; (f) Tracing results drawn from ST with a high threshold (purple) and a low threshold (yellow), respectively, and from ST-LFV (red). The manual tracing results are labeled with green curves. The location of the weak neurite in (b) and (c) is labeled with circles.

Furthermore, we showed that ST-LFV is superior to SparseTracer when tracing neurites with 12 experimental image stacks. Each sub-block was 600 × 600 × 600 voxels. Two typical datasets and their corresponding tracing results are provided (Figs. 6a and b). These two datasets are extracted from the whole brain dataset, and their sites are labeled with arrows in Fig. 6c. The sites of ten other datasets are also given (Fig. 6c). These sites show that these twelve datasets are distributed among different brain regions. When using SparseTracer to analyze these datasets, we set the tracing threshold value to maximize the average tracing accuracy. This is why, for some datasets with strong backgrounds, as shown in the middle of Fig. 6a, SparseTracer confuses noise with the neurite’s signals and produces more tracing results than ST-LFV. We quantified the tracing results from SparseTracer and ST-LFV (Figs. 6d and e). The average recall rate and the average precision rate are 0.93 and 0.86 for SparseTracer, and 0.99 and 0.97 for ST-LFV, respectively. The formula used to calculate the recall and precision rates can be found in our previous work (T. Quan et al. 2016).

**Figure 6.**
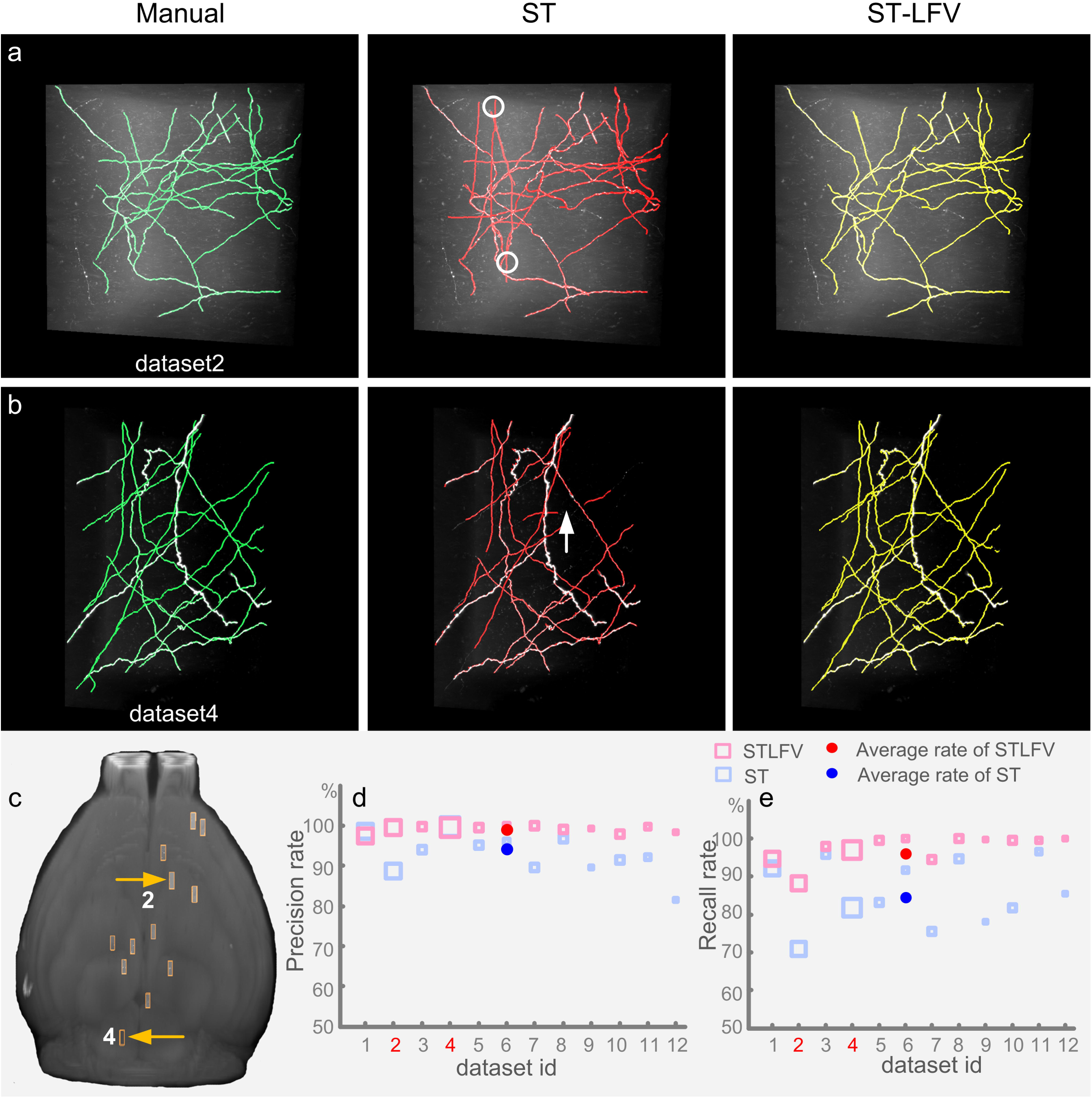
Comparisons of neurite tracing results drawn from SparseTracer (ST) and ST-LFV. (a) One typical dataset with a strong background and its corresponding tracing results drawn manually (green), with ST (red), and with ST-LFV (yellow). Over-tracing results from ST are labeled with circles; (b) Another dataset with a weak background in which ST fails to trace part of a neurite labeled with an arrow; (c) The distribution of the datasets’ locations in the mouse brain. The locations of dataset (a) and (b) are labeled with yellow arrows; (d) and (e) are automatic tracing results from the datasets in (c) and are measured with a precision rate (d) and a recall rate (e), respectively. The red numbers '2'&'4' on the lateral axis represent the datasets shown in (a) and (b).

Our premises are the basis of ST-LFV. If the features of neuronal image stacks are consistent with our premises, then ST-LFV can effectively identify neurites with weak signals from these image stacks and provide perfect tracing results. Here, we examined whether our premises apply in general to DIADEM datasets. Two datasets were used for this purpose, both of which are labeled by green fluorescent protein. One of the datasets is shown in Fig. 7a (Neocortical Layer 6 Axons dataset) and was imaged by a two-photon microscope, and the other (OP dataset) was imaged by 2-channel confocal microscopy. Using the same procedure described in the Methods section, we extracted foreground points and background points and calculated corresponding feature vectors. The calculated feature vectors (Figs. 7c-f) demonstrate large differences between the extracted foreground and extracted background points. These differences result in an identification model for detecting weak signals with an extremely low training error (0%/2.6% for Fig. 7g and 0%/0% for Fig. 7h). These results indicate that along with the MOST datasets, our premises hold for the DIADEM dataset. Note that the extracted background points also include a few points located in regions adjacent to or at the boundaries of neurites, especially for images (Fig. 7a) in which the number of signal voxels is not extremely small compared to the total number of voxels. In this case, these few points can be regarded as positive for the training classifier (see Fig. 7g).

**Figure 7.**
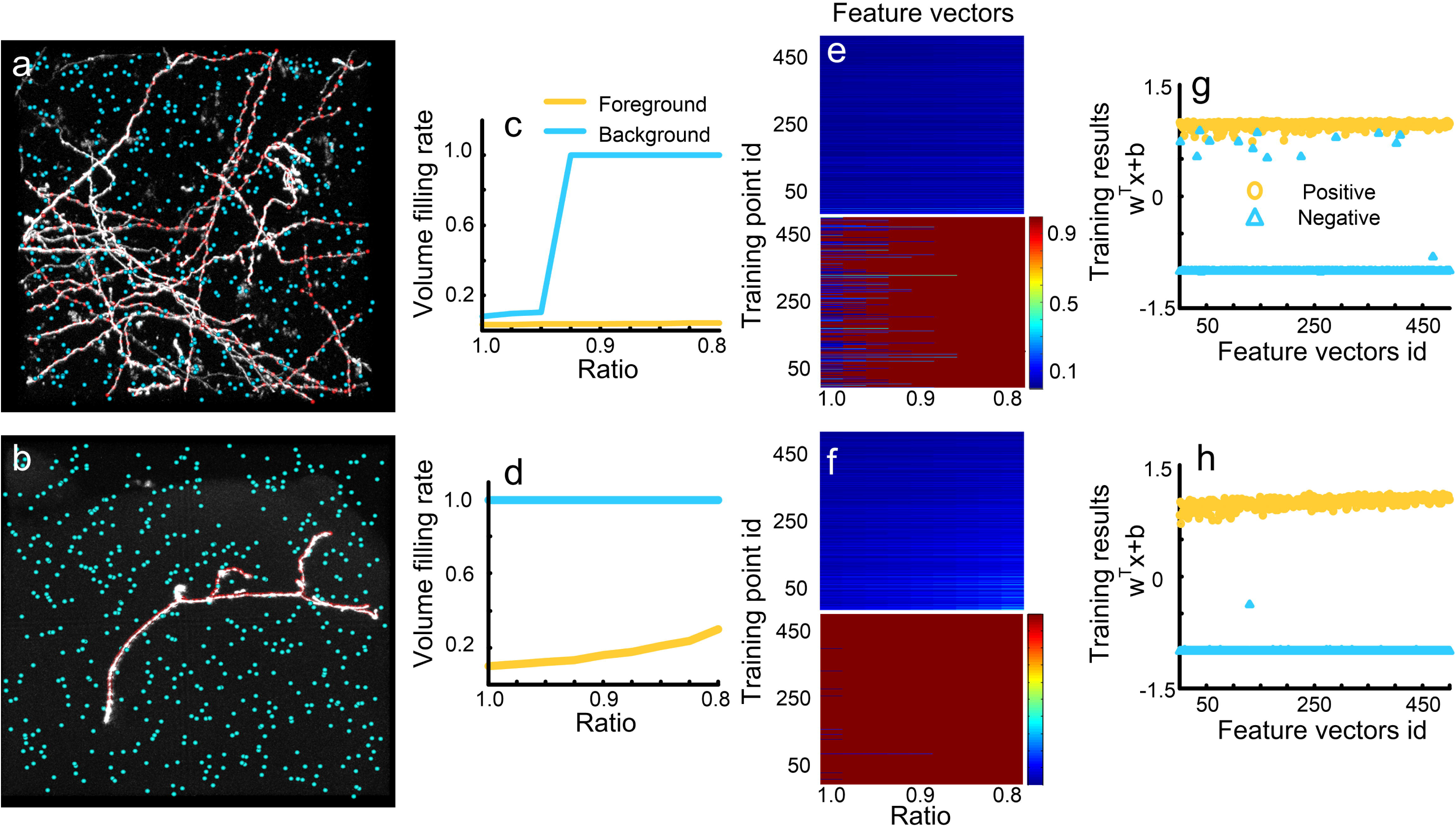
Feature vector extraction from DIADEM datasets. (a) and (b) are two datasets, one from the Neocortical Layer 6 Axons datasets and the other from the Olfactory Projection Fibers datasets. The foreground (red) and background (blue) voxels used for feature extraction; (c) and (d) The foreground (yellow) and background (blue) feature vectors calculated from two labeled voxels in (a) and (b), respectively; (e) and (f) are feature vectors of all labeled foreground (upper) and background (bottom) voxels in (a) and (b), respectively; (g) and (h) generate the SVM identification model with feature vectors from (e) and (f), respectively. The positive (yellow circles) and the negative (blue triangles) results correspond to foreground and background feature vectors.

We also demonstrated that our premises hold for BigNeuron datasets. Two typical datasets (checked6_mouse_tufts and checked_mouse_korea) were used for this purpose (Figs. 8a and b). Both datasets have strong background noise, but the respective feature vectors for the extracted foreground and background points are different and are easily classified into two groups (Figs. 8c-f). Furthermore, these feature vectors, as the training set, can be used to derive identification models with a training error rate of 0.4%/0.4% for Fig. 8g and 0.8%/1.6% for Fig. 8h, respectively. The low training errors indicate the large differences between the features of the foreground and background points. These differences originate from the fact that our premises are consistent with the features of the analyzed datasets. Note that in Figs. 8g and h, some training errors occur that can be attributed to the interference of the strong background.

**Figure 8.**
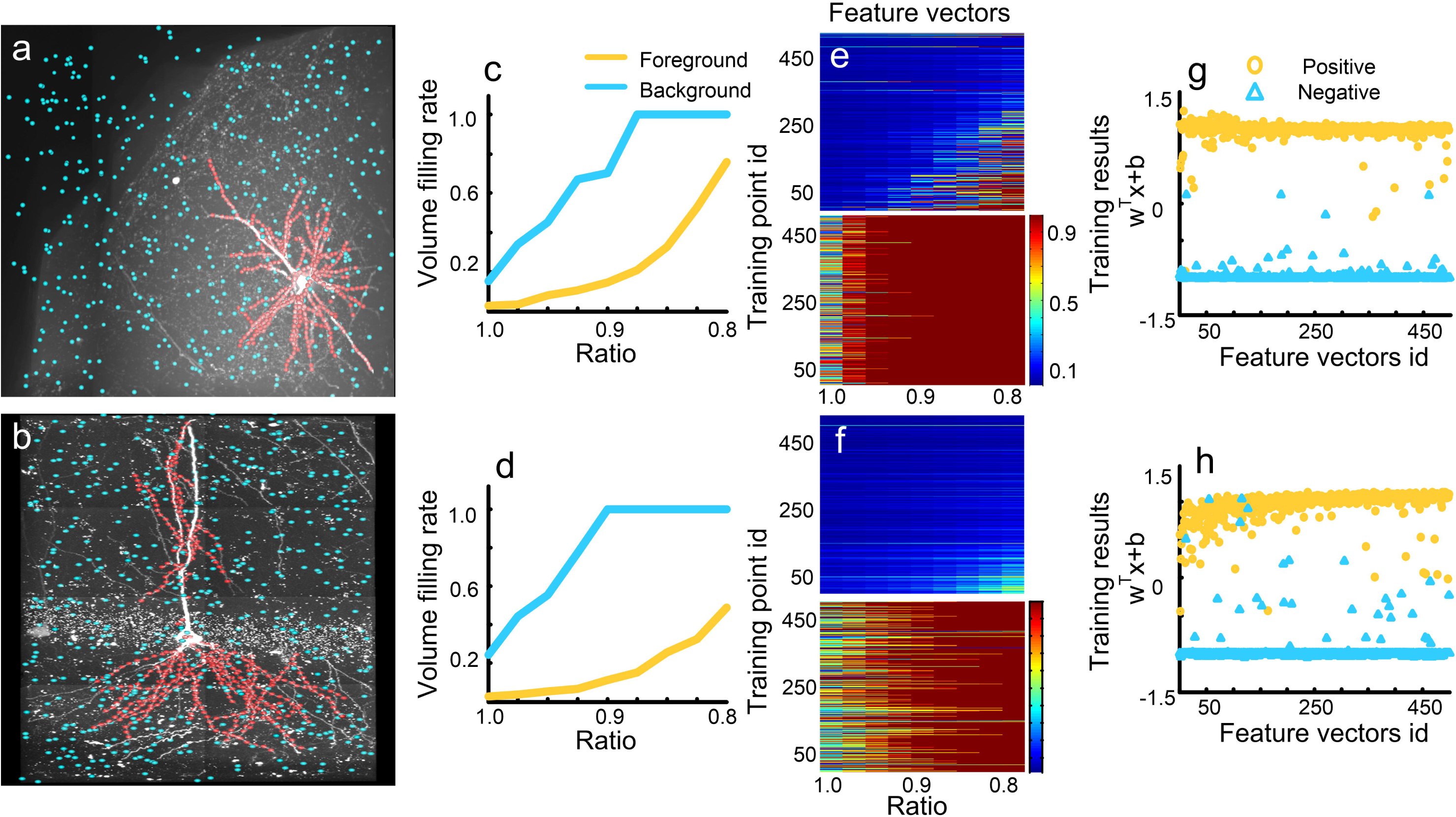
Feature vector extraction from BigNeuron datasets. (a) and (b) are two datasets, one from the checked6_mouse_tufts datasets and the other from the checked_mouse_korea datasets. The foreground (red) and background (blue) voxels used for feature extraction; (c) and (d) The foreground (yellow) and background (blue) feature vectors are calculated from two labeled voxels in (a) and (b), respectively; (e) and (f) are feature vectors of all labeled foreground (upper) and background (bottom) voxels in (a) and (b), respectively; (g) and (h) generate the SVM identification model with feature vectors from (e) and (f), respectively.

We demonstrated that ST-LFV can address the challenges illustrated in Fig. 8. The two datasets selected include weak signals and strong background noise. We compared the tracing results from SparseTracer and ST-LFV (Figs. 9a-d), and found that SparseTracer cannot trace some neurites with extremely weak signals (arrows in Fig. 9a and c). ST-LFV provided tracing results that included almost all of the neurites that could not be traced with SparseTracer. In addition, this result verified that the identification model (Figs. 8g and h) is usable even with some training errors.

**Figure 9.**
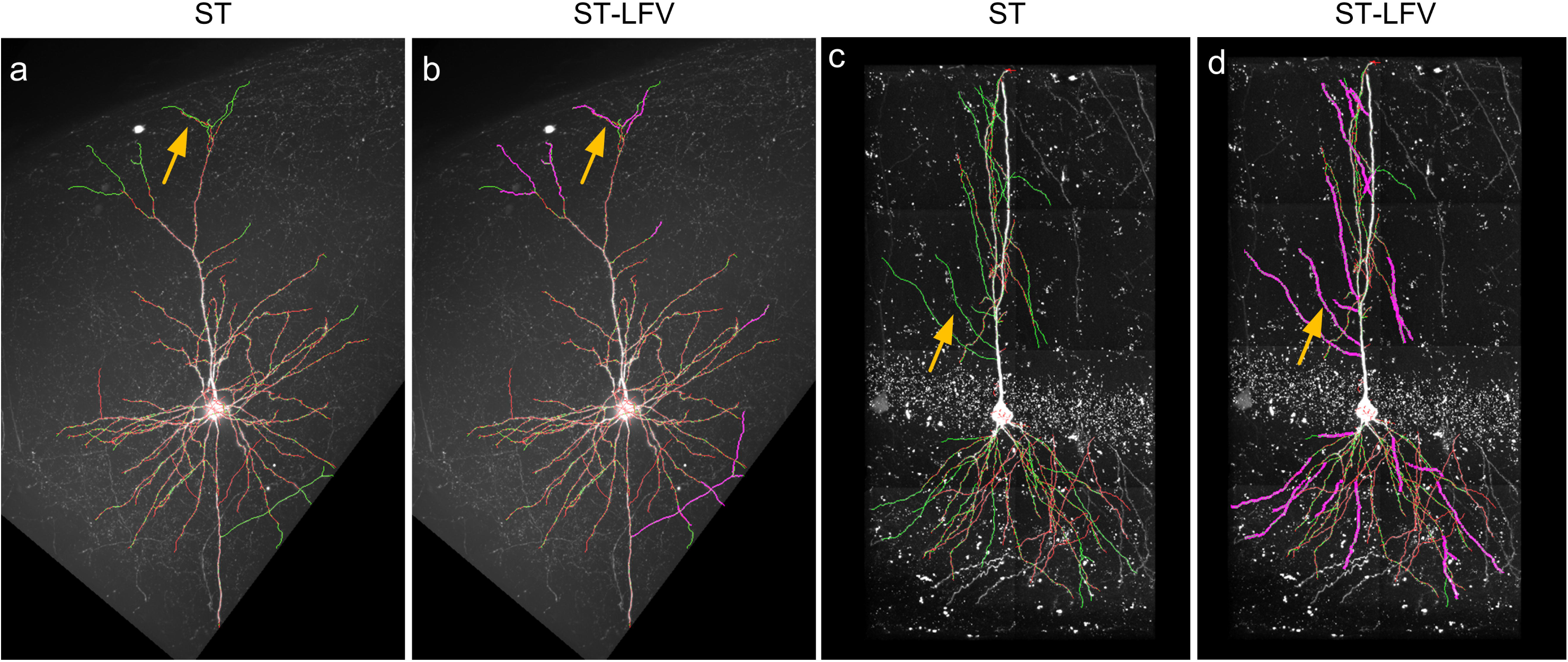
Tracing performance on two BigNeuron datasets drawn from SparseTracer (ST) and ST-LFV. (a) The tracing results drawn from ST (red). Some neurites cannot be detected by ST, and these are labeled with arrows. The ground truth (green) is also provided. (b) ST-LFV can provide tracing results (purple) that are almost equivalent to the ground truth (green). The neurites that cannot be detected with ST can be traced successfully (see the labeled arrow); (c) and (d) share descriptions similar to (a) and (b), respectively.

Due to the superior tracing performance of ST-LFV, we found that it could vastly enhance the automation level of SparseTracer software, enabling the rapid tracing of neurites at a large scale. To illustrate this property, we selected a dataset that included several long axons, with a total size of 1.99 × 1.93 × 1.32 mm^3^ (voxel size, 0.3 × 0.3 × 1 μm^3^, 105 GB). We adopted the divide-and-conquer strategy used in our previous work (S. Li et al. 2016) for large dataset analysis. We also added a manual editing module into the SparseTracer tool, which helps to obtain an initial tracing direction and sites for continuous tracing when neurites with weak signals are not detected by the algorithm. Due to the manual editing module, SparseTracer and ST-LFV can provide the same tracing results (Figs. 10b and c). However, the number of manual editing sessions required for SparseTracer (20 times) is far greater than for ST-LFV (1 time). This shows the advantage of ST-LFV in large-scale tracing of neurites. We quantified this advantage by comparing the total time required by ST-LFV with that of SparseTracer. In this tracing, ST-LFV required approximately 5 minutes, and SparseTracer required approximately 20 minutes. In ST-LFV, the time required to identify weak signals is 47% of the total time, which is less than the time required for tracing neurites. Using SparseTracer to trace neurites requires intensive manual editing and to provide a fair comparison we counted the total time that a skillful annotator spent on this tracing. Therefore, 20 minutes is a conservative estimate for SparseTracer. Image reading time is not included in the total time, as a RAID is connected to the workstation directly and the time required can be neglected here. Using the above comparisons, we conclude that ST-LFV significantly improves the tracing performance of SparseTracer and is valuable for large-scale tracing of neurites.

**Figure 10.**
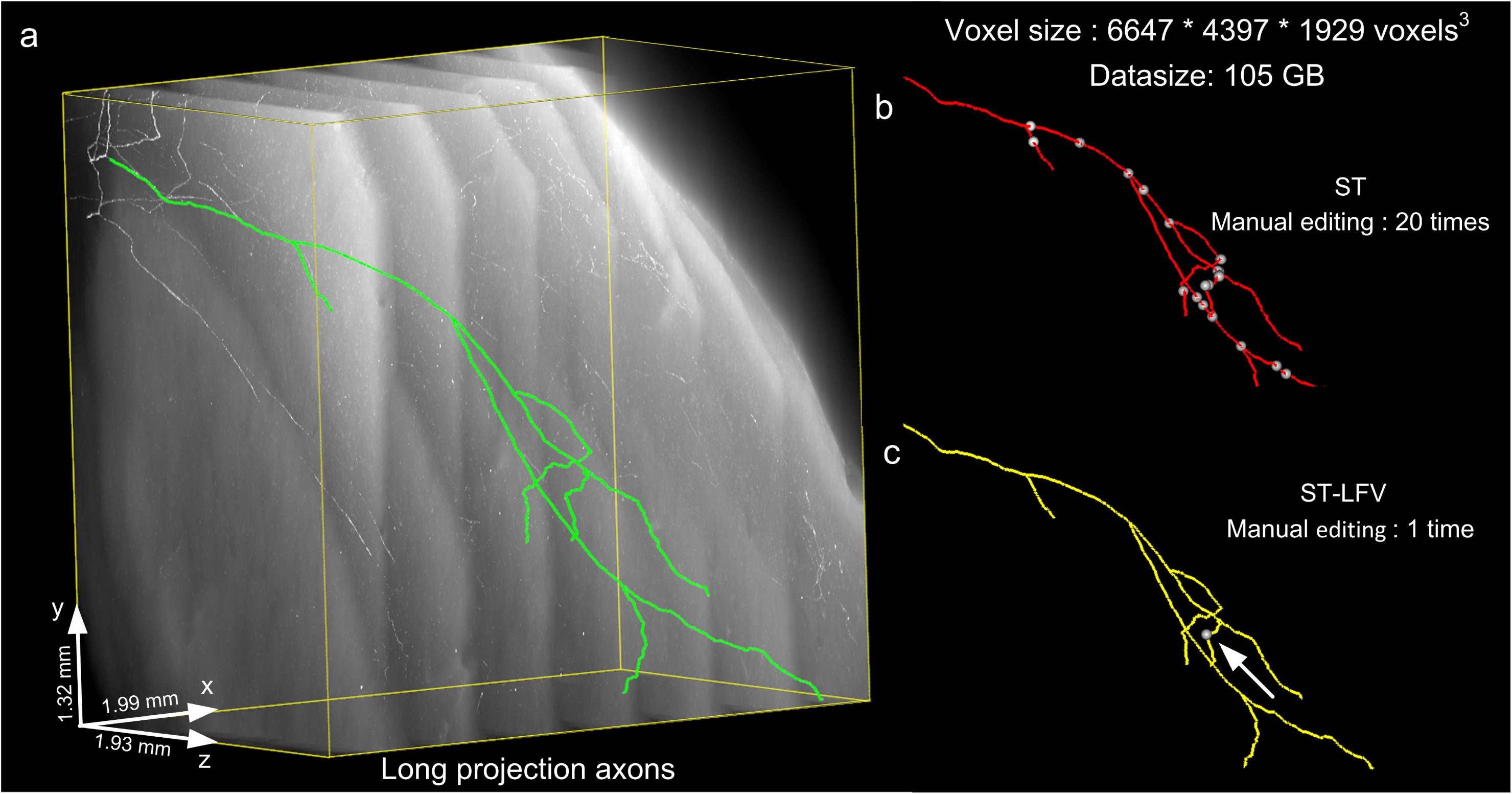
Tracing neurites at a large scale. (a) A large-scale dataset (approximately 105 GB) and the tracing results (ground truth) drawn from a human annotator (green); (b) the tracing results (red) drawn from SparseTracer (ST) equivalent to the ground truth. A total of 20 manual edits (interferences) are required, and their corresponding locations are labeled with white dots; (c) for ST-LFV, only one manual edit is required (arrow).

## 4. Discussion

ST-LFV involves more rules deduced from images by human beings than other methods (R. Li et al. 2017; Chen et al. 2015). These rules are based on the fact that our premises are commonly applicable to most neuronal images collected with optical microscopes, and provide a basis for constructing a feature vector that displays the differences between foreground and background voxels. Using more rules avoids a complicated procedure for extracting valid features and identifying weak signals, indicating that intensive computation is avoidable. This is the primary reason why ST-LFV is suitable for large-scale tracing of neurites.

In ST-LFV, the identification model is embedded into the tracing procedure, which is different from other methods (R. Li et al. 2017; Chen et al. 2015). When the tracing termination conditions are effective, the identification method works and identifies whether the current tracing point is a foreground voxel. If yes, tracing is continuous. The identification model enhances the ability of ST-LFV to trace neurites with weak signals and is linked to the tracing procedure. Other methods separate the identification and tracing procedures by using a machine learning method to identify as many foreground voxels as possible and then perform the tracing procedure, i.e., extract the skeleton of the identified foreground. These methods aim to identify all foreground voxels, and thus have the relatively high computational complexities, which is an obstacle to large-scale tracing of neurites. ST-LFV activates the identification model only when the tracing terminal conditions are effective, and thus only identifies a few foreground voxels when tracing a neurite. This contributes to the ability of ST-LFV to rapidly trace neurites in large-scale images.

Similar to machine learning methods (Suykens and Vandewalle 1999; Cortes and Vapnik 1995), obtaining a training set is a key part of ST-LFV. Here, the training set contains positive (foreground) and negative (background) feature vectors. In our method, the training set is automatically generated and does not require manual labeling. A positive feature vector in the training set is determined by its corresponding foreground voxel (see the Methods section). The voxel is automatically generated by the tracing procedure, such as SparseTracer (S. Li et al. 2016) or other methods (Bas and Erdogmus 2011; Rodriguez et al. 2009). A negative feature vector is determined by its corresponding voxel randomly (from a uniform distribution) chosen from the image. This random selection assures that the chosen voxel has an extremely low probability of being in the foreground. This low probability is ensured by the fact that the spatial distribution of neurites is sparse and the number of foreground voxels is extremely small compared to the total number of voxels. According to the above analysis, automatically obtaining a training set is feasible.

In SparseTracer, we use a constrained principal curve to trace neurites with weak signals (S. Li et al. 2016; T. Quan et al. 2016). In general, it is difficult to obtain a forward tracing direction from this type of neurite because of inadequate local structure information. In this case, the constrained principal curve introduces directional information for the traced points, and this direction becomes the forward tracing direction. This feature allows SparseTracer to detect weaker signals compared to other methods (Rodriguez et al. 2009; Wang et al. 2011; Xiao and Peng 2013). When analyzing images that include weak signals, SparseTracer demonstrates highly accurate tracing performance (>85% recall at >90% precision). However, SparseTracer, like most other methods, uses a set of thresholds to determine whether a tracing termination situation has been reached. This termination condition may not be suitable for the detection of weak signals from an inhomogeneous background (See Figs. 4g and h). This type of weak signal detection is a common task when tracing large-scale neurites. Considering this situation, we proposed an identification model and combined it with SparseTracer, i.e., ST-LFV, for large-scale tracing of neurites. To provide better tracing performance, our proposed identification method can be embedded into the pipeline of most tracing methods, such as the voxel scoping method (Rodriguez et al. 2009), the model fitting method (Zhao et al. 2011), the principal curves method (Bas and Erdogmus 2011), and others.

The identification model used in ST-LFV is based on rules drawn from various types of neuronal images. The identification method can be used directly for most neuronal images. However, in a few cases, some processing may be necessary before using the identification method. For example, if the neuronal images contain densely distributed somas, a certain amount of foreground voxels will appear in the randomly chosen voxels that are used to construct the negative training set. To some extent, this violates the assumption (see the Methods section) that the probability that the randomly chosen voxels contain some foreground voxels is extremely small.

In this case, by identifying the soma region as in our previous work (Tingwei Quan et al. 2013; T. Quan et al. 2014), voxels in the soma region can be removed from the negative training set. For images with densely distributed neurites, the difficulties in obtaining the negative training set and the corresponding solutions are similar to those in images with dense somas. For images with local backgrounds that are not smooth, some de-noising methods (Meijering 2010) can be used to generate a smooth background.

## 5. Conclusion

We proposed a method to identify neurites with weak signals from an inhomogeneous background. We verified that the extracted feature, which differentiates the foreground and background, is widely applicable across various types of datasets. We demonstrated that the identification method can vastly increase the accuracy of tracing neurites. We further demonstrated that this identification method is suitable for large-scale tracing of neurites, which aids in the reconstruction of a neuron across different brain regions or even the whole brain.

## Acknowledgments

This work was supported by the Science Fund for Creative Research Group of China (Grant No. 61421064), National Natural Science Foundation of China (Grant No. 81327802), National Program on Key Basic Research Project of China (Grant No. 2015CB7556003) and Director Fund of WNLO.

